# BCL2L13 Influences Autophagy and Ceramide Metabolism without Affecting Temozolomide Resistance in Glioblastoma

**DOI:** 10.1101/2024.08.23.609447

**Authors:** Courtney Clark, Amir Barzegar-Behrooz, Simone C. da Sila Rosa, Kianoosh Naghibzadeh, Mehdi Eshraghi, Jaodi Jacobs, Xiaohui Weng, Abhay Srivastava, Rui Vitorino, Sudharsan Rao Ande, Amir Ravandi, Sanjiv Dhingra, Stevan Pecic, Donald Miller, Shahla Shojaei, Saeid Ghavami

## Abstract

Temozolomide (TMZ) resistance in glioblastoma (GBM) arises through metabolic rewiring that links mitochondrial function, autophagy balance, and sphingolipid metabolism. TMZ resistant (R) U251 cells exhibited suppressed apoptosis and complete blockade of autophagy flux, evidenced by LC3II and p62 accumulation and insensitivity to Bafilomycin A1. BCL2L13, strongly upregulated in R cells, emerged as a dual regulator of mitophagy and ceramide metabolism. BCL2L13 knockdown (KD) produced opposite effects in TMZ sensitive (NR) and resistant cells: in NR cells, KD elevated LC3II, reduced respiratory reserve, and triggered compensatory lipid synthesis; in R cells, KD decreased LC3II without restoring flux or TMZ sensitivity. Lipidomic profiling revealed that BCL2L13 loss reactivated CerS6 in NR cells, increasing C16:0 and mid-chain ceramides, while relieving CerS2 inhibition in R cells, elevating very long chain (C22 to C24) and glycosylated ceramides. These distinct sphingolipid signatures were confirmed by PLS-DA and KEGG enrichment, which highlighted steroid hormone, arachidonic, and linoleic acid metabolism in NR KD cells versus neuroactive ligand-receptor and signaling pathways in R KD cells. Together, these findings position BCL2L13 as a molecular integrator of mitochondrial respiration, autophagy flux, and CerS-dependent lipid remodeling, unveiling a context-specific metabolic mechanism that supports GBM cell survival under chemotherapeutic stress.

## INTRODUCTION

Glioblastoma (GB) is the most aggressive and lethal primary brain tumor, with a median survival of only 15 months despite aggressive multimodal treatment [1-3]. One of the significant challenges in treating GB is the development of resistance to temozolomide (TMZ), the standard chemotherapy for this disease [4, 5]. The underlying mechanisms driving TMZ resistance (R) involve complex cellular processes, including alterations in autophagy, apoptosis, and lipid metabolism [6, 7]. Understanding these processes is crucial for developing strategies to overcome resistance and improve therapeutic outcomes.

Autophagy is a critical cellular process that involves the degradation and recycling of cellular components to maintain homeostasis. Autophagy plays a dual role in cancer, acting as a tumor suppressor and a survival mechanism [8, 9]. In R cells, our recent findings have shown that autophagy flux is inhibited, as evidenced by the accumulation of autophagy markers such as LC3βII and SQSTM1 [5]. This inhibition suggests a blockage in the autophagic process, potentially allowing these cells to evade apoptosis—a process typically activated in response to cellular stress and damage induced by chemotherapy [5]. These results indicate that the disruption of autophagy flux in R cells is a critical factor in developing resistance, as it may enable the cells to withstand the cytotoxic effects of TMZ.

BCL2L13, a member of the BCL-2 family of proteins, has been identified as a crucial regulator of apoptosis and ceramide synthesis in cancer cells [10]. In GB, BCL2L13 is highly expressed and plays a significant role in regulating the synthesis of ceramides, a class of lipids involved in regulation apoptosis [11]. Our study further reveals that BCL2L13 interacts with ceramide synthases CerS2 and CerS6, changing their activity and thus affecting the balance of ceramides within the cell. This modulation of ceramide levels by BCL2L13 contributes to the survival of GB cells under TMZ treatment, as ceramides are known to induce apoptotic pathways that can lead to cell death. Given the role of BCL2L13 in regulating autophagy flux and ceramide synthesis [11-14], we hypothesize that BCL2L13 contributes to TMZ resistance in GB by controlling the fate of these pathways. Specifically, we propose that the elevated expression of BCL2L13 in R cells disrupts autophagy flux and alters ceramide metabolism, leading to enhanced survival of resistant cells. This study aims to elucidate the mechanistic roles of BCL2L13 in R cells, focusing on its interactions with autophagy, ceramide pathways, and ceramide synthases and provide opportunity to understand role of BCL2L13 in developing TMZ chemoresistance in GB.

To test this hypothesis, we conducted a detailed analysis of autophagy flux, apoptosis, and ceramide profiles in R and NR cells with varying levels of BCL2L13 expression. Our objectives were to determine how BCL2L13 influences autophagy, ceramide synthesis, and apoptosis in the context of TMZ resistance and to assess whether targeting BCL2L13 could restore sensitivity to TMZ and improve therapeutic outcomes in GB. The findings from this study provide new insights into the molecular mechanisms underlying TMZ resistance and suggest novel therapeutic targets to enhance the effectiveness of GB treatment.

## MATERIALS AND METHODS

### Cell Culture

All experiments used the U251-mKate human glioblastoma cell line (developed by Dr. Marcel Bally, Experimental Therapeutics, British Columbia Cancer Agency, Vancouver, BC, Canada). Cells were cultured in Dulbecco’s Modified Eagle’s Medium-high glucose (4 mg/ml) (DMEM) (CORNING; Cat #: 50-003-PB) supplemented with 10% Fetal Bovine Serum (FBS) (Gibco™; Cat #: 16000044) and 1% penicillin and streptomycin. Cells were maintained in a humidified incubator in standard cell culture conditions (95% air and 5% CO_2_ at 37°C). Stable BCL2L13 knockdown (KD) and negative control (scramble) were grown in the same medium plus puromycin (4 μg/ml) to ensure the selection of stably transfected cells.

### Preparation of Resistant Cells

U251-mKate cells were cultured in T75 tissue culture flasks in a complete media containing high glucose DMEM, 10% FBS (Gibco; Cat #:10437-036), and 1% Penicillin-Streptomycin (Gibco, Cat #:15140122) and incubated at 37°C, 5% CO_2_, in a humidified incubator (Thermo Fischer Scientific, US). A pulsed-selection strategy was chosen to make a clinically relevant chemoresistant model [15]. Once cells reached 70% confluence, they were treated with 100μM TMZ (Sigma Aldrich, CAS #: 85622-93-1) over three weeks, followed by four weeks of TMZ-free media for recovery. Cells were cultured in complete media with 250μM TMZ for three weeks, followed by another four weeks with TMZ-free media. The final population of U251-mKate cells resistant to 250μM TMZ was selected and cultured in a complete media without TMZ for four weeks.

### Immunoblotting

Protein expression was detected using western blotting, with GAPDH and Ponceau S as loading controls. Cells were lysed in buffer (20 mM Tris-HCl pH 7.5, 0.5 mM PMSF, 0.5% Nonidet P-40, 100 μM β-glycerol 3-phosphate, 0.5% protease and phosphatase inhibitor cocktail), and lysates were centrifuged at 10,000 g for 10 min at 4°C. Protein concentration was determined by Lowry assay. Proteins were separated by SDS-PAGE and transferred to PVDF membranes. After blocking with 5% skim milk in TBST for 1 hour at room temperature, membranes were incubated overnight at 4°C with primary antibodies in 1% skim milk/TBST, followed by HRP-conjugated secondary antibodies for 2 hrs at room temperature. Detection was performed using ECL (Amersham-Pharmacia Biotech) and visualized with a BioRad imager. Band intensity was quantified using Alpha Ease FC software [16, 17].

### Transmission Electron Microscopy (TEM)

Autophagy was confirmed using TEM. Briefly, cells were cultured in 100mm dishes in DMEM (10% FBS 1% P/S) in standard cell culture incubator conditions. Cells were fixed in Karnovsky fixative for 1 hour at 4°C, and the pellet was resuspended in 5% sucrose in 0.1M Sorenson’s phosphate buffer overnight at 4°C. Cells were pelleted and post-fixed with 1% osmium tetroxide in 0.1M Sorenson’s buffer for 2 hours. Next, cells were dehydrated and embedded in Embed 812 resin for sectioning. Thin sections (200 nm) were cut and placed on copper grids for staining and counter-staining with uranyl acetate and lead nitrate, respectively. Imaging was done using a Philips CM10 electron microscope [17].

### Immunocytochemistry (ICC)

Changes in autophagy in WT cells and BCL2L13 knockdown transfectants were detected using immunocytochemistry. Cells were fixed with 3.7% formaldehyde for 30 mins at room temperature, washed thrice with DPBS, and permeabilized overnight at 4°C in blocking buffer (5% IgG-free BSA, 0.3% Triton X-100 in DPBS). The blocking buffer was then replaced with antibody solution (1% BSA, 0.3% Triton X-100, and conjugated antibodies in DPBS), and samples were incubated overnight at 4°C in the dark. Anti-LC3 primary antibody (1:2000, Sigma) and donkey anti-rabbit IgG secondary antibody (1:2000, Jackson ImmunoResearch) were used. After staining, samples were washed, incubated with DPBS, and stored at 4°C in the dark until imaging. Imaging was performed using a Zeiss Axiovert 200 microscope with Calibri 7 LED Light Source and Axiocam 702 mono camera. Image processing and quantification were done using Fiji (ImageJ) and Zen 2.3 Pro software. All experiments were independently repeated 3 times, with a minimum of 10 images captured per condition [17].

### MTT Viability Assay

Cell viability of U251 GB cells was measured using the MTT assay. Cells were cultured to 50% confluence in high glucose DMEM with 10% FBS and 1% P/S, then seeded (NR: 50,000 cells/mL; R: 75,000 cells/mL) in 96-well plates. Cells were treated with varying concentrations of BafA1 (1, 5, or 10nM) for 72h. MTT (20μL of 5mg/mL) was added to each well, and plates were incubated at 37°C with 5% CO2 for 3 hrs. The media was aspirated, and 200μL of DMSO was added to dissolve the formazan crystals. After 25 mins incubation at RT in the dark, absorbance was measured at 570nm using a Synergy H1 Microplate Reader. Relative cell viability was calculated as the percentage of control using the formula: (mean OD570 of treated cells/mean OD570 of control cells) x 100 [18].

### Production of Stable BCL2L13 Knock Down (KD) Cell Line

U251 R (R) and NR GB (NR) cells were seeded in 12-well plates and cultured in DMEM (10% FBS, 1% P/S). At 40% confluency, cells were treated with 10 µg/mL polybrene in DMEM (without FBS) for 1 hour, followed by infection with lentiviruses carrying shRNA targeting BCL2L13 or scramble RNA (scRNA) as a control. After 12 hours of incubation, media were refreshed, and cells were allowed to recover for 48 hours. The transfection process was repeated to enhance efficiency. Transfected cells were selected with 4 µg/mL puromycin. Successful knockdown of BCL2L13 was confirmed by western blotting in stable clones.

### Flow Cytometry/Cell Cycle Analysis

U251 GB cells were seeded in 12-well plates (50,000 cells/well for NR and 75,000 cells/well for R cells) and grown to 60% confluence. Cells were treated with 250 µM TMZ for 48 or 96 hours. Post-treatment, cells were detached with EDTA buffer, harvested by centrifugation, and washed in PBS. They were resuspended in hypotonic PI lysis buffer (0.01% Triton X-100, 1% sodium citrate, 40 µg/mL propidium iodide) and incubated at room temperature in the dark for 30 minutes. Flow cytometry was performed using a Beckman Coulter Cytoflex LX with a 488 nm laser and 610/20 BandPass filter to measure the percentage of cells in Sub-G0, G0-G1, S, and G2-M phases [18].

### Measurement of Mitochondrial Respiration: Sea Horse Assay

Oxygen consumption rate (OCR) was assessed using the Agilent Seahorse XFe24 analyzer. U251 GB cells (NR (scramble), R (scramble), NR-BCL2L13-KD, and R-BCL2L13-KD) were seeded in XF 24-well microplates at 0.3 x 10□ cells/well. On the experiment day, cells were switched to XF base minimal DMEM with glucose, L-glutamine, and sodium pyruvate, then incubated at 37°C for 1 hour. Baseline OCR was measured, followed by protein leak (1 µM oligomycin), maximal respiration (2 µM FCCP), and non-mitochondrial respiration (1 µM rotenone, 1 µM antimycin A). OCR normalization was performed using live cell counts stained with Hoechst33342. Maximal respiration was calculated as FCCP OCR minus non-mitochondrial OCR; spare capacity as FCCP OCR minus baseline OCR; ATP production as baseline OCR minus Oligomycin OCR; and proton leak as Oligomycin OCR minus non-mitochondrial OCR. Data are presented as pmol O□/min/cell count.

### Co-Immunoprecipitation (Co-IP) for BCL2L13 Interaction with CerS2 and CerS6 in U251 R and NR Cells

R and NR cells were lysed in buffer (20 mM Tris-HCl, pH 7.5, 150 mM NaCl, 1% NP-40, 0.5% sodium deoxycholate, 1 mM EDTA, 1 mM PMSF, 10% glycerol, protease/phosphatase inhibitors). Lysates were pre-cleared with Protein A/G agarose beads, then incubated overnight at 4°C with anti-BCL2L13 antibody, followed by a 2-hour incubation with fresh beads. Immunocomplexes were washed, eluted in 2X SDS-PAGE sample buffer, and analyzed via Western blot using antibodies against CerS2 and CerS6. Densitometry was performed to quantify interactions.

### Phase Contrast

Bright-field images were taken using an Axio Observer ZEISS microscope (ZEISS, Oberkochen, Germany) using 20x and 50x magnification objectives. Quantification, scale bars, background subtraction, and processing were done on Fiji (ImageJ) and Zen 2.3 Pro software.

### Liquid Chromatography-Mass Spectrometry (LC-MS) for Lipidomics Analysis of Ceramide Species

Lipids were extracted from U251 GB cells (NR, R, NR-BCL2L13-KD, R-BCL2L13-KD) using a chloroform (2:1, v/v) mixture. Cells grown to confluence were collected, sonicated, and centrifuged (1000g, 10 min). The supernatant was mixed with the internal standard (ISTD), vortexed, and centrifuged (3500 rpm, 5 min). The lower phase was dried under N□ gas and reconstituted in water-saturated butanol, followed by methanol with 10 mM ammonium formate. After centrifugation (10000g, 10 min), lipids were analyzed by LC-MS. Separation was performed on a Zorbax C18 column using a linear gradient of mobile phases A and B, both containing 10 mM ammonium formate. The elution program involved increasing from 0% to 100% mobile phase B over 8 minutes, with re-equilibration to starting conditions before the next injection. Diacylglycerol and triacylglycerol species were separated isocratically at 100 μl/min. The column was maintained at 50°C, and the injection volume was 5 μl.

### Mass Spectral Analysis

Lipids eluted from the HPLC system were introduced into the AbSciex 4000 QTRAP triple quadrupole linear hybrid mass spectrometer. The mass spectrometer was operated in scheduled Multiple Reaction Monitoring (MRM) mode. 322 unique lipids spanning 25 lipid classes/subclasses were screened for targeted semi-quantitation. All lipid species other than fatty acids were scanned in positive electrospray ionization mode [ESI+]. The individual lipids in each lipid class were identified by lipid class-specific precursor ions or neutral losses. Lipids were then quantified by comparing the deisotoped lipid peak areas against those of the class-specific ISTDs added before lipid extraction. The total carbon number of the fatty acids represented lipids. In ESI+ mode, the instrument settings were optimized as follows: curtain gas (psi), 26; collision gas (nitrogen), medium; ion spray voltage (V), 5500; temperature (°C), 500.0; ion source gas 1 (psi), 40.0; ion and source gas 2 (psi), 30.0. The MRM detection window was fixed between 45s and 90s depending upon the chromatographic peak width of the lipid class. Isobaric species within the same class, such as PC(O) and PC(P), exhibited clear separation in this method. Also, molecular species within the same lipid class, which differ only in the number of double bonds, were well separated chromatographically. All analyses were performed with R software (version 4.1.1). Limma, pheatmap, and ggplot2 packages were used.

### SMILES Code Retrieval and Annotation

SMILES (Simplified Molecular Input Line Entry System) codes for lipid species identified in the LC–MS lipidomics analysis were retrieved from the Human Metabolome Database (HMDB; https://hmdb.ca). For each statistically significant lipid, the corresponding SMILES code was cross-validated with the LC–MS fragmentation data to ensure that the molecular structure matched the experimentally determined head group, acyl chain composition, and degree of unsaturation. When multiple structural variants were available, the canonical SMILES format was selected to ensure consistency. Verified SMILES codes were used for downstream cheminformatics analyses, including pathway identification.

### Statistics and reproducibility

All experiments were conducted in triplicate, with three independent repeats. Data are presented as mean ± standard deviation. Statistical significance was determined using a two-tailed t-test or one-way/two-way ANOVA with Tukey’s post hoc test for multiple comparisons, considering p-values ≤ 0.05 as significant. Lipid classification and annotation were performed using the LIPIDMAPS database. Lipidomic data were analyzed with MetaboAnalyst 5.0, including hierarchical clustering, principal component analysis (PCA), and partial least squares discriminant analysis (PLSDA) to identify differential lipids.

## RESULTS

### Temozolomide induces more cell death in U251 Temozolomide non-resistant GB cells

We found that R cells maintained 50% viability at TMZ concentrations up to 500μM (*P* < 0.0001), compared to significant cell death in NR cells (Fig. 1A). R cells were maintained in 250μM TMZ, with the drug removed 24 hours before experiment. Phase contrast microscopy revealed that NR cells showed reduced confluence and increased cell death after 48 hours of treatment with 100μM TMZ, while R cells remained unaffected (Fig. 2B). R cells were smaller, rounder, and had shorter filopodia; in contrast to the elongated filopodia of NR cells. Nicolletti apoptosis assay confirmed that TMZ significantly increased the sub-G0 apoptotic phase (*P* < 0.0001) and altered cell cycle distribution in NR cells, with no significant effects on R cells (Fig. 1C-E). No significant changes were observed in the S phase for either cell type following treatment (*P* > 0.05) (Fig. 1F).

**Figure 1.**
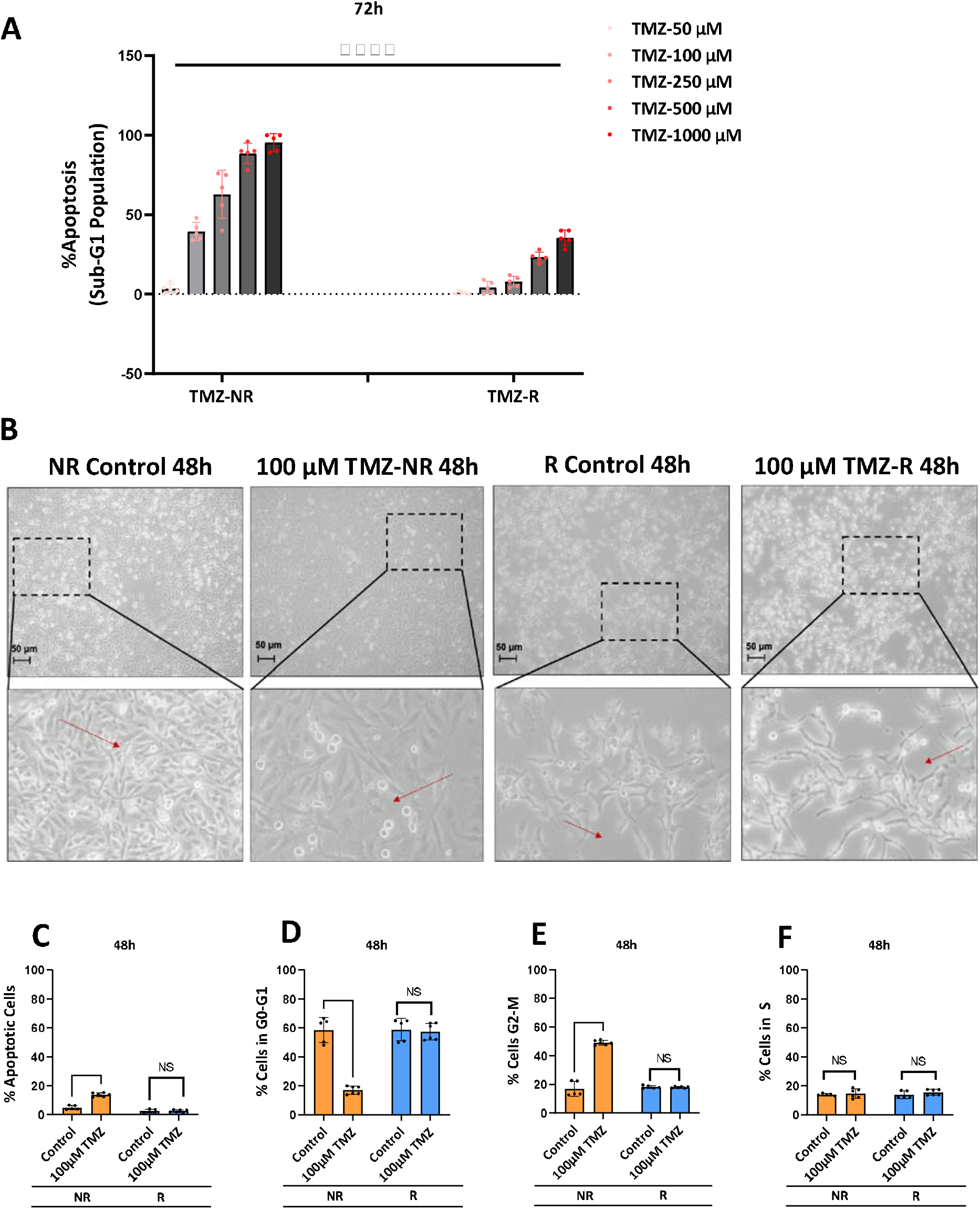
Temozolomide induces more cell death in U251 Temozolomide NR cells. **A**. R and NR cells were treated with 10–5000 μM TMZ, and apoptosis was assessed over 72 hours using the Nicolleti assay. **B** Phase-contrast images of R and NR cells after 48-hour treatment with 100 μM TMZ show more apoptotic morphology in NR cells. **C-F** TMZ (100 μM) treatment induced significantly higher apoptosis and increased G2/M phase arrest in NR cells compared to R cells, which showed no significant change. Data represent mean ± SD of three independent experiments.

### Autophagy flux is inhibited in TMZ-Chemoresistant GB cells

We investigated autophagic activity in R and NR cells by assessing LC3βII and SQSTM1 (p62) expression levels using immunoblotting. LC3βII and SQSTM1, key autophagy markers, were degraded over 72 hours in NR cells, indicating active autophagy. In contrast, their accumulation in R cells suggested a significant disruption in autophagy flux (*P* < 0.01) (Fig. 2A, B). To further examine autophagy, we used Bafilomycin A1 (BafA1), an inhibitor of vacuolar H (+)-ATPase that blocks autophagy flux. MTT assays demonstrated that BafA1 (10 nM) significantly reduced the viability of NR cells (*P* < 0.001) but had no cytotoxic effect on R cells (Fig. 2C). Subsequently, we selected 5 nM BafA1 to compare autophagy flux between R and NR cells. In NR cells, BafA1 treatment increased LC3βII and SQSTM1 levels (*P* < 0.01), whereas in R cells, BafA1 had no significant impact, suggesting complete disruption of autophagy at the autophagosome-lysosome fusion stage (Fig. 2D, E). Immunocytochemistry (ICC) confirmed these findings, showing more LC3β puncta in R cells, indicative of accumulated autophagosomes (Fig. 2F). Additionally, transmission electron microscopy (TEM) revealed a significantly higher number of double-membrane vacuoles in R cells compared to NR cells, further supporting the conclusion that autophagy is disrupted in R cells (Fig. 2G, H).

**Figure 2.**
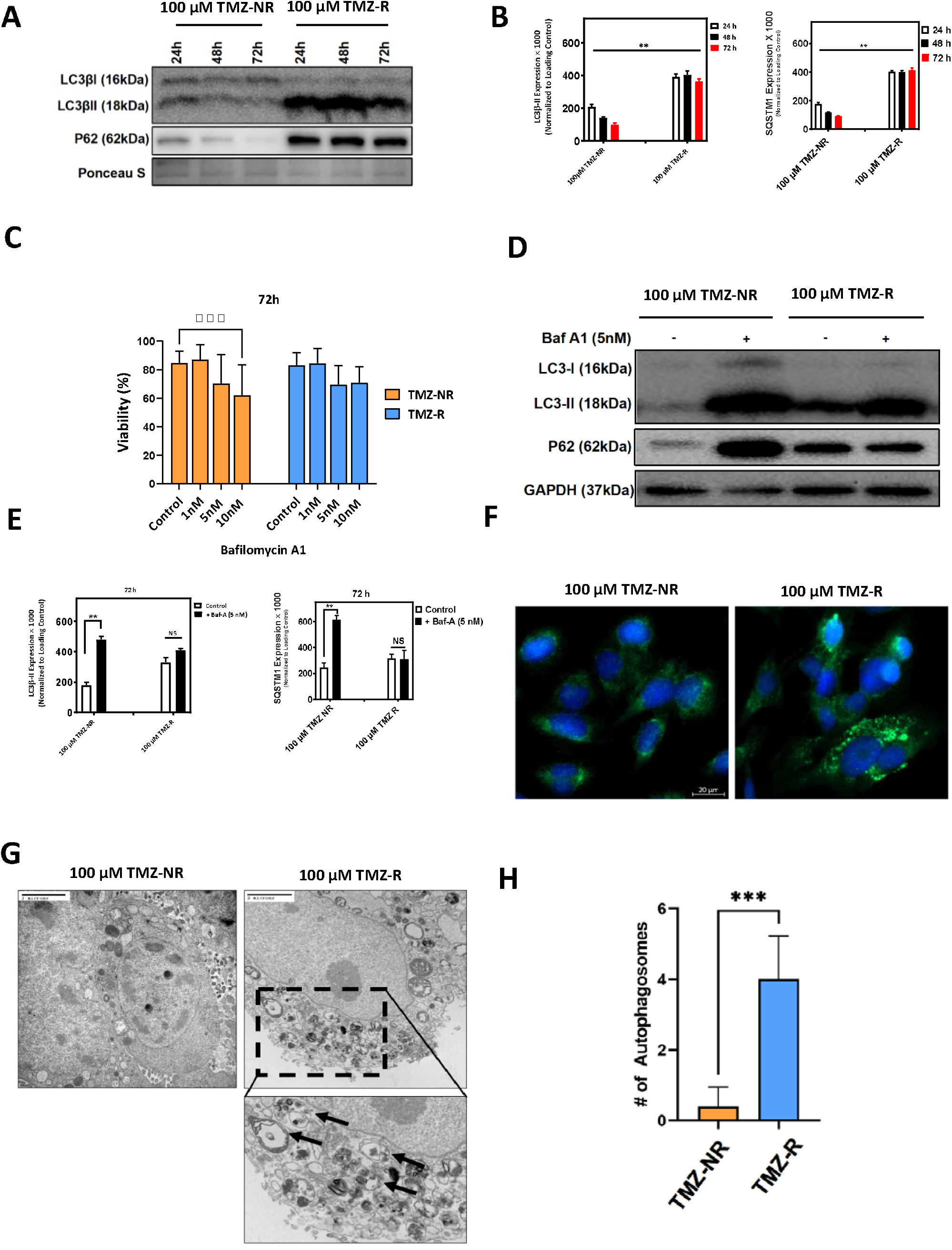
Autophagy flux is inhibited in R cells. **A, B**. NR and R cells were cultured for 72 hours with lysates collected every 24 hours for immunoblotting. Autophagy markers (SQSTM1 degradation, LC3β lipidation, and LC3βII formation) were assessed, showing active autophagy flux in NR cells over time, but inhibition in R cells (P < 0.01). Data represent three biological replicates. **C**. Cytotoxic effects of Bafilomycin-A1 (1, 5, 10nM) on NR and R cells over 72 hours were measured by MTT assay, revealing significant cell death induction in NR cells at 10nM (P < 0.001), with minimal effect on R cells. **D, E** Immunoblot analysis of NR and R cells treated with 5nM Bafilomycin-A1 for 72 hours, with lysates collected every 24 hours, demonstrated inhibited autophagy flux in NR cells (P < 0.01) but no effect in R cells. GAPDH was the loading control. **F** Immunocytochemistry for LC3β puncta showed increased autophagosome formation in R cells relative to NR cells. **G, H** TEM images (25,000x) of NR and R cells demonstrated significantly more double-membrane autophagic vacuoles in R cells (P < 0.001), with quantification across 10 cells.

### BCL2L13 Orchestrates Resistance-Linked Autophagy Modulation Without Affecting Apoptosis in Glioma Cells

BCL2L13 expression was examined in R and NR cells using immunoblotting, with samples collected every 24 hours over 72 hours. The results demonstrated significantly higher BCL2L13 expression in R cells compared to NR cells at the 48 and 72-hour time points (*P* < 0.01) (Fig. 3A, B). Given the known roles of BCL2L13 in apoptosis regulation and mitochondrial function, we further explored its impact on autophagy and cell survival in R cells. To investigate the effects of BCL2L13 on autophagy, apoptosis, and TMZ response, BCL2L13 was knocked down in U251 R and NR cells via lentiviral transfection, achieving stable RNA silencing (Fig. 4C). Transmission electron microscopy (TEM) revealed that BCL2L13 knockdown (KD) did not significantly alter the number of cytosolic double-membrane vacuoles in either NR or R cells (*P* > 0.05) (Fig. 3C, D). Autophagy flux was further assessed using Bafilomycin A1 (Baf-A1, 5 nM, 72 hours). In NR cells, BCL2L13 KD significantly increased LC3β-II levels, while in R cells, it significantly decreased LC3β-II without Baf-A1 treatment (Fig. 3E, F). Additionally, BCL2L13 KD did not significantly affect SQSTM1 degradation in NR cells but significantly reduced SQSTM1 degradation in R cells (Fig. 3E, F). Baf-A1 treatment significantly increased LC3β-II and decreased SQSTM1 degradation in NR cells, irrespective of BCL2L13 expression, while not affecting these markers in R cells (Fig. 3G, H). These findings suggest that BCL2L13 is key in regulating autophagy flux, particularly in R cells where autophagy is inhibited. Finally, we evaluated the impact of BCL2L13 expression on TMZ-induced apoptosis and cell cycle distribution using flow cytometry. BCL2L13 expression did not significantly alter apoptosis in either NR or R cells after 48 and 96 hours of TMZ treatment (*P* > 0.05). However, BCL2L13 KD in NR cells led to a significant reduction in the percentage of cells in the G2-M phase after 48 hours, while R cells (both scramble and BCL2L13 KD) showed a decrease in the G0-G1 phase after 96 hours of treatment (Fig. 3H, I).

**Figure 3.**
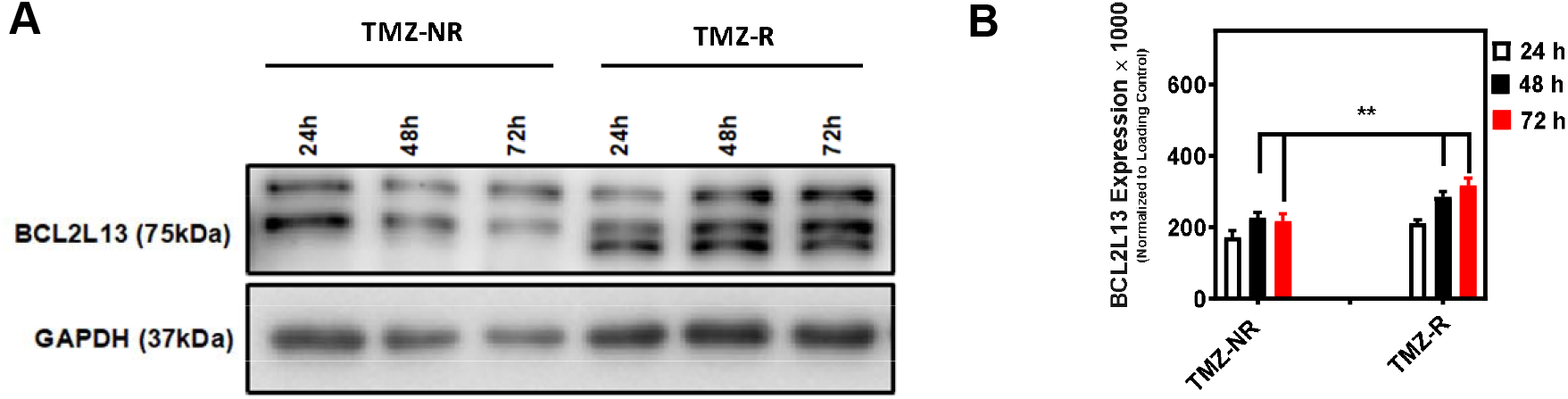
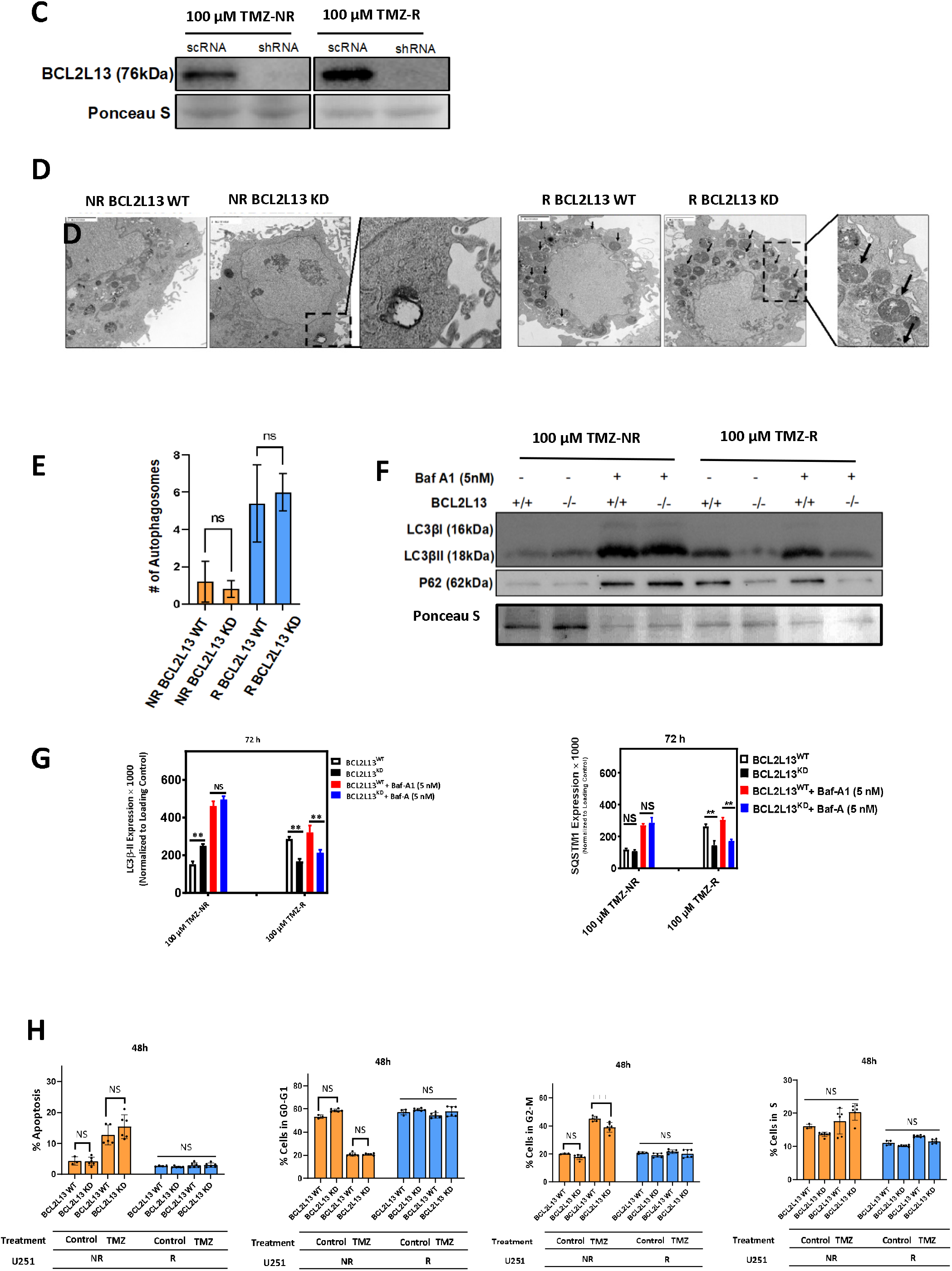
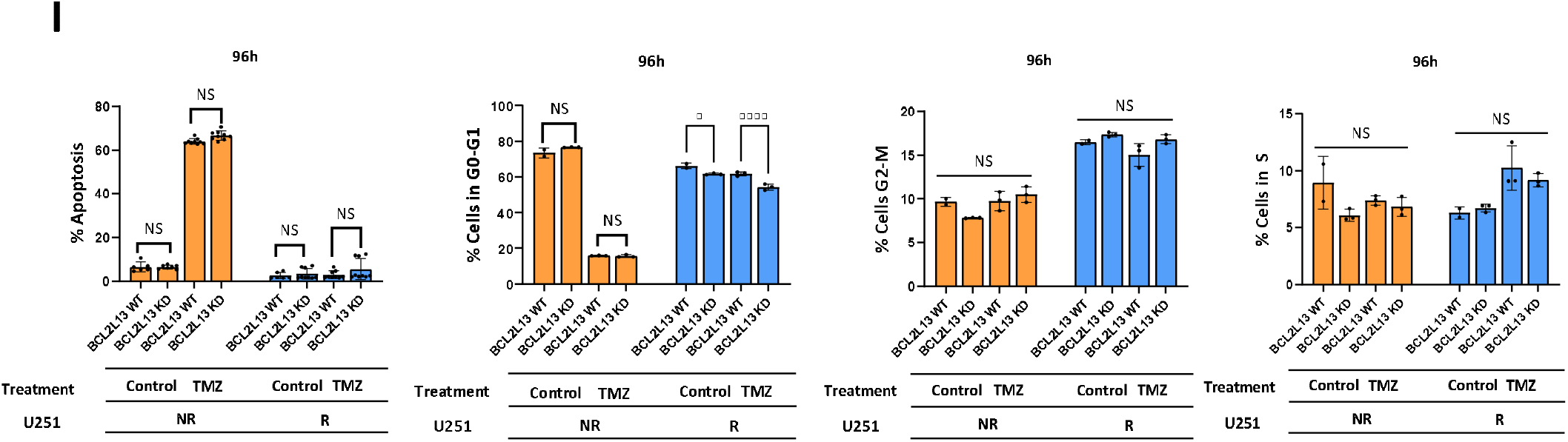
BCL2L13 is Upregulated in TMZ-Resistant Cells and Modulates Autophagy Without Affecting Apoptosis. (A, B) Immunoblotting revealed significantly elevated BCL2L13 in U251 R vs. NR cells at 48 and 72 hours (P < 0.01). GAPDH served as a loading control. (C) Lentiviral knockdown of BCL2L13 in NR and R cells was confirmed by immunoblot. (D, E) TEM imaging showed no significant change in autophagosome number after BCL2L13 knockdown in either cell type (P > 0.05). (F, G) BCL2L13 knockdown altered LC3β-II and p62 levels in a cell context-dependent manner. In NR cells, LC3β-II increased (P < 0.01) without affecting p62; in R cells, LC3β-II decreased and p62 degradation increased (P < 0.01). Bafilomycin-A1 impaired autophagy flux in NR but not R cells. (H, I) Flow cytometry showed no significant effect of BCL2L13 knockdown on TMZ-induced apoptosis or cell cycle distribution (P > 0.05). Data represent n = 3 (biological) ± SD.

### BCL2L13 Maintains Mitochondrial Respiratory Reserve and Integrity in TMZ-Sensitive Glioblastoma Cells

Previous studies have established the involvement of BCL2 family proteins in various aspects of mitochondrial function [19]. Figure 5 illustrates the effects of BCL2L13 knockdown (KD) on mitochondrial respiration in TMZ-sensitive (NR) and -resistant (R) glioblastoma cells. In Panel A, OCR traces reveal that NR control cells maintain the highest mitochondrial activity throughout, whereas R KD cells show a marked drop in FCCP-induced respiration, indicating impaired mitochondrial flexibility. Panel B confirms significantly lower basal respiration in R control versus NR control cells (p□<□0.05), with no further reduction upon BCL2L13 KD, suggesting BCL2L13 is dispensable for resting metabolism. Panel C shows that maximal respiratory capacity is high in NR controls but significantly reduced in R and R KD cells (p□<□0.0001); BCL2L13 KD modestly reduces capacity in NR cells, mirroring the resistant phenotype, yet has no additive effect in R cells. Panel D reveals that spare respiratory capacity is significantly greater in NR controls than all other groups (p□<□0.0001), indicating that both TMZ resistance and BCL2L13 loss independently compromise bioenergetic reserve. Panel E shows that ATP-linked respiration is highest in NR controls, significantly reduced in R and R KD cells (p□<□0.001), and modestly decreased in NR KD cells, which still outperform R KD (p□<□0.05), underscoring BCL2L13’s role in ATP production. Proton leak, shown in Panel F, is elevated in all KD and R conditions (p□<□0.0001), with the highest levels in R KD cells, pointing to mitochondrial membrane dysfunction. Panel G demonstrates significantly higher non-mitochondrial oxygen consumption in NR KD than R KD cells (p□<□0.05), suggesting metabolic compensation unique to the sensitive background. Overall, BCL2L13 sustains mitochondrial respiratory capacity, ATP production, and membrane integrity in TMZ-sensitive GBM cells, while its loss recapitulates features of resistant bioenergetic collapse. In contrast, BCL2L13 depletion does not further impair already-compromised mitochondrial function in resistant cells, highlighting its selective importance in maintaining metabolic adaptability in sensitive states.

**Figure 4.**
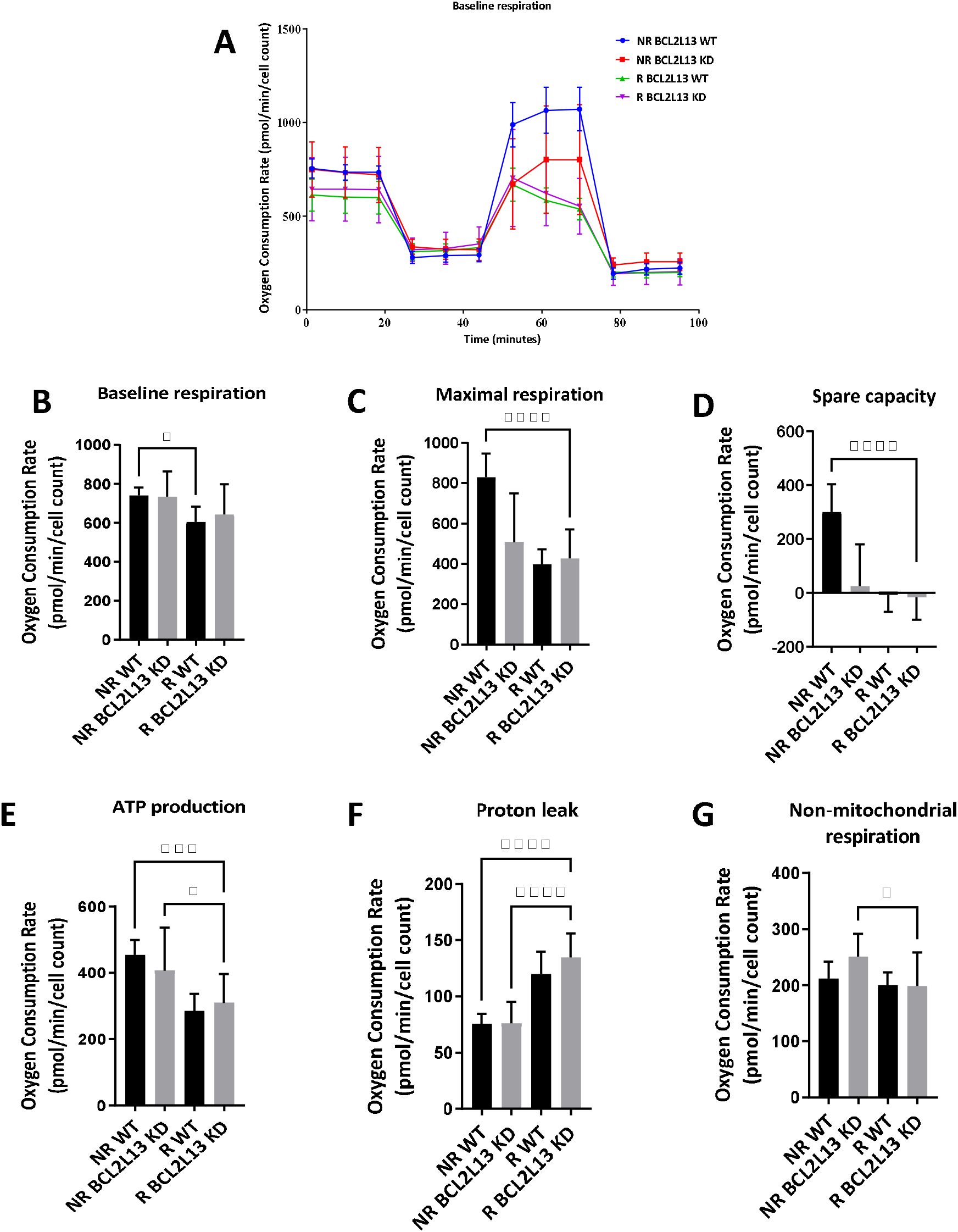
BCL2L13 Preserves Mitochondrial Function and Energetic Flexibility in TMZ-Sensitive Glioblastoma Cells. Figure 5 shows the effects of BCL2L13 knockdown (KD) on mitochondrial respiration in U251 NR and R glioblastoma cells. (A) OCR profiles indicate that NR control cells maintain robust respiration, while R KD cells exhibit the sharpest decline following FCCP treatment, reflecting impaired mitochondrial flexibility. (B) Basal respiration is significantly lower in R control than NR control cells (p□<□0.05), with no further reduction upon BCL2L13 KD. (C) Maximal respiration is significantly diminished in R control and R KD cells compared to NR controls (p□<□0.0001), and modestly reduced in NR KD cells. (D) Spare respiratory capacity is highest in NR control cells and significantly reduced in all other groups (p□<□0.0001). (E) ATP-linked respiration follows a similar pattern, with NR KD cells showing an intermediate phenotype between NR and R KD (p□<□0.001). (F) Proton leak is significantly increased in all KD and R groups (p□<□0.0001), especially in R KD cells. (G) Non-mitochondrial OCR is significantly higher in NR KD compared to R KD (p□<□0.05), indicating greater compensatory metabolic activity in TMZ-sensitive cells. All data represent n = 5 biological replicates and are shown as mean ± SD.

### TMZ Resistance Alters BCL2L13–Ceramide Synthase 2 and 6 Interactions in Glioblastoma Cells

Previous studies have identified physical interactions between BCL2L13 and ceramide synthases 2 and 6 and its effect of regulation of their activity in glioblastoma, implicating these complexes in the regulation of mitochondrial membrane composition and apoptosis signaling. Building on this, we aimed to determine whether TMZ resistance modifies the interaction landscape between BCL2L13 and CerS2 or Ceramide CerS6—enzymes responsible for generating distinct ceramide species. This investigation was prompted by our earlier findings showing that BCL2L13 supports mitochondrial respiratory capacity predominantly in NR cells while affecting autophagy flux in R cells, suggesting a context-dependent regulatory role. Using co-immunoprecipitation, we found that BCL2L13 exhibited stronger binding to CerS6 in NR cells, while its interaction with CerS2 was enhanced in R cells (Fig. 5A, B). This resistance-associated shift in binding preference suggests that BCL2L13 may differentially modulate ceramide metabolism depending on treatment status—potentially contributing to the altered mitochondrial integrity and metabolic inflexibility observed in R cells. These results further position BCL2L13 as a molecular integrator of lipid metabolism and mitochondrial stress adaptation in glioblastoma.

**Figure 5.**
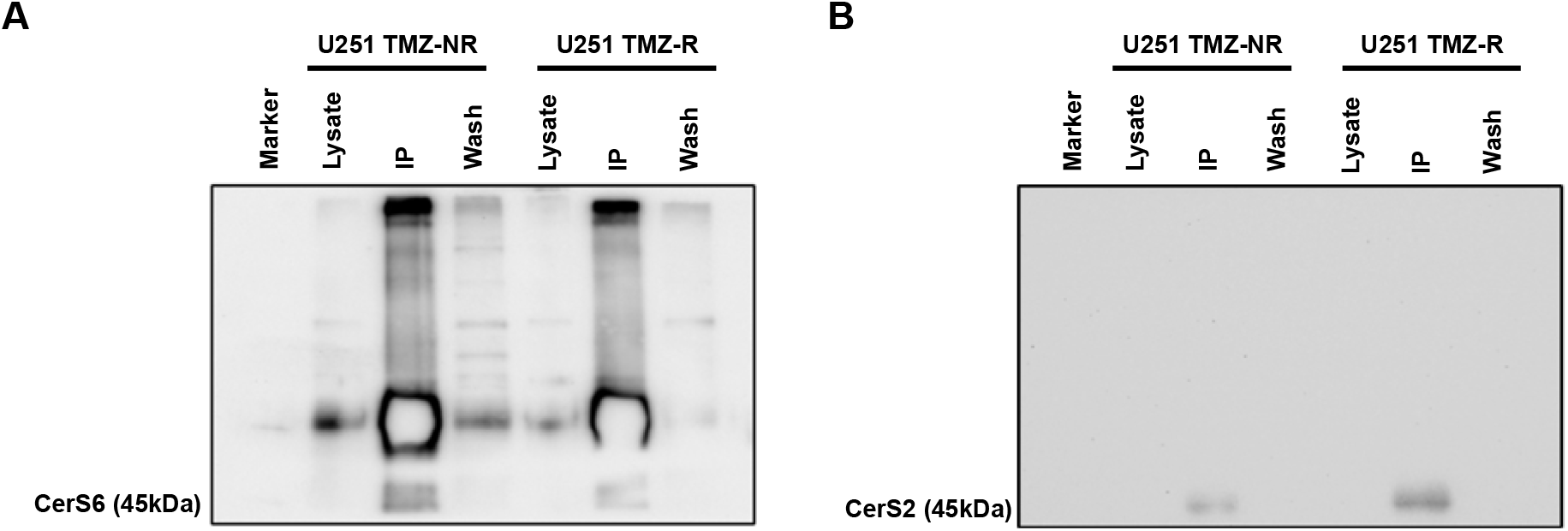
BCL2L13 interacts differently with Ceramide Synthase 2 and 6 in NR and R cells. (A,B) U251 NR and U251 R cells were cultured for 72 hours, and cell lysates were collected for co-immunoprecipitation to determine CerS6 and BCL2L13 interaction. The results showed that CerS6 interacted more with BCL2L13 in U251 NR cells than in U251 R cells. B U251 NR and U251 R cells were cultured for 72 hours, and cell lysates were collected for co-immunoprecipitation to determine CerS2 and BCL2L13 interaction. Our results showed that CerS2 had lower interaction with BCL2L13 in U251 NR cells than in U251 R cells.

### BCL2L13 Knockdown Induces Resistance-Specific Remodeling of Ceramide Profiles and Lipid Signaling via CerS2 and CerS6 Deregulation

Building on our co-immunoprecipitation findings that BCL2L13 binds and inhibits CerS6 in non-resistant (NR) cells and CerS2 in resistant (R) cells (Fig. 5A, B), we hypothesized that BCL2L13 knockdown (KD) would relieve this inhibition and reshape the ceramide landscape accordingly. Because CerS6 primarily generates C14–C16 ceramides and CerS2 synthesizes very-long-chain (VLC) ceramides (≥ C22), we analyzed lipid species across multiple ceramide subclasses. Heatmap analysis (Fig. 6A) revealed that both NR and R scramble controls exhibit higher levels of dihydroceramides (dhCer) and monohexosylceramides (MHC) compared to their KD counterparts, indicating that wild-type BCL2L13 limits CerS activity. In R KD cells—where BCL2L13–CerS2 binding is reduced—we observed increased levels of dihexosylceramides (DHC) and trihexosylceramides (THC) with VLC fatty acyl chains (e.g., 18:0, 20:0, 22:0), consistent with enhanced CerS2 activity. Notably, reductions in THC 16:0 and THC 24:0/24:1 in R KD cells reflect both expected CerS6 activation and potentially altered downstream glycosylation flux. These nuanced lipid shifts suggest that BCL2L13 modulates both synthesis and post-synthetic remodeling of chain-specific sphingolipids in a resistance-dependent manner. Principal Component Analysis (PCA) further confirmed distinct ceramide signatures between TMZ-sensitive and resistant states (Fig. 6B).

Detailed analysis of individual ceramide species (Fig. 6C–G) corroborated this regulatory model. Cer 16:0 was most enriched in R KD and lowest in NR KD, matching CerS6 reactivation upon BCL2L13 loss. VLC ceramides such as Cer 18:0–22:0 were elevated in R WT vs. NR WT but declined in R KD, again suggesting BCL2L13 modulates CerS2 output. Cer 24:0 and 24:1 showed complex regulation, with peak levels in R WT and inconsistent shifts post-KD, pointing to resistance-specific lipid processing mechanisms. dhCer profiles (Fig. 6D) showed sustained VLC enrichment in R lines, especially R WT, with dhCer 16:0 dropping upon KD in R cells. MHC and DHC subclasses (Fig. 6E–F) mirrored these patterns, with highest levels in R WT and reductions following BCL2L13 KD. THC species (Fig. 6G) showed greater variability: THC 18:0–22:0 were elevated in R KD, while THC 24:0 and 24:1 varied across groups, suggesting differential downstream handling of ceramides beyond direct CerS activity. Collectively, these subclass-specific and chain-length–specific profiles reinforce BCL2L13’s role in shaping sphingolipid output through selective inhibition of CerS2 and CerS6, with unique consequences in TMZ-sensitive versus resistant backgrounds.

To pinpoint lipid species most impacted by BCL2L13 loss, we performed PLS-DA. In NR cells, four lipids—Cer 20:0, DHC 20:0, DHC 24:1, and DHC 24:0—emerged as key discriminators (VIP ≥ 1.4), marking a robust BCL2L13-dependent signature (Fig. 6H). These matched our earlier prediction that loss of CerS6 inhibition in NR KD would increase medium and VLC ceramides. In R cells, PLS-DA identified Cer 22:0 and THC 18:0 as discriminative species (Fig. 6I), with Cer 22:0 elevation directly aligning with increased CerS2 activity after BCL2L13 KD. THC 18:0, though not a CerS2 product, may reflect glycosylation flux downstream of elevated VLC ceramide synthesis. To contextualize these lipidomic shifts within broader signaling, KEGG pathway enrichment was performed. In NR KD cells, five FDR-significant pathways were identified—including steroid hormone biosynthesis, linoleic and arachidonic acid metabolism—highlighting the metabolic breadth of BCL2L13 deficiency (Fig. 6J). In contrast, R KD cells showed strong enrichment of neuroactive ligand–receptor interaction and calcium signaling pathways (Fig. 6K), indicating distinct post-lipid remodeling outcomes shaped by resistance. Together, these findings demonstrate that BCL2L13 governs ceramide enzyme output and downstream lipid remodeling in a resistance-specific manner, integrating metabolic control and sphingolipid signaling under chemotherapeutic stress.

**Figure 6.**
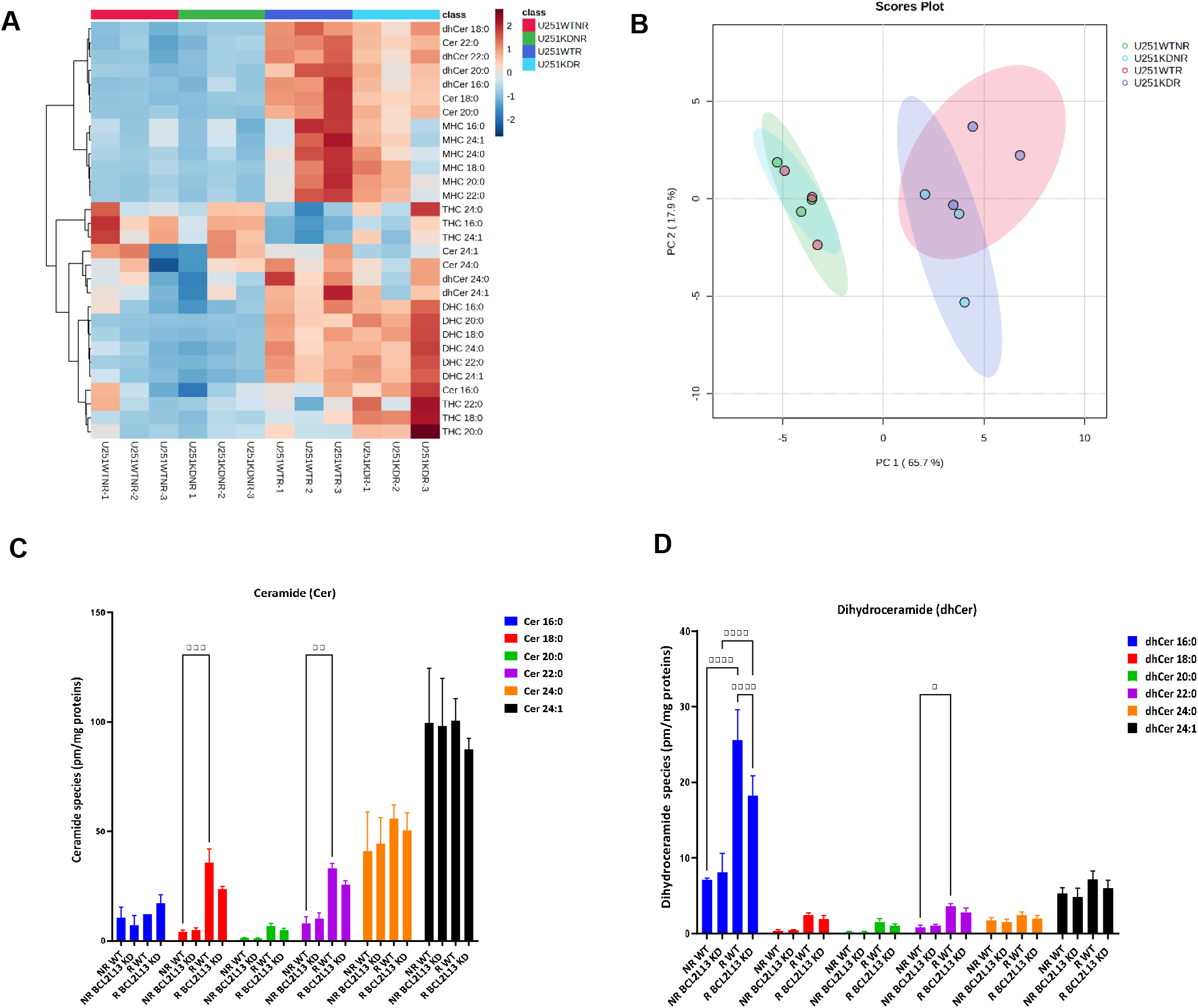
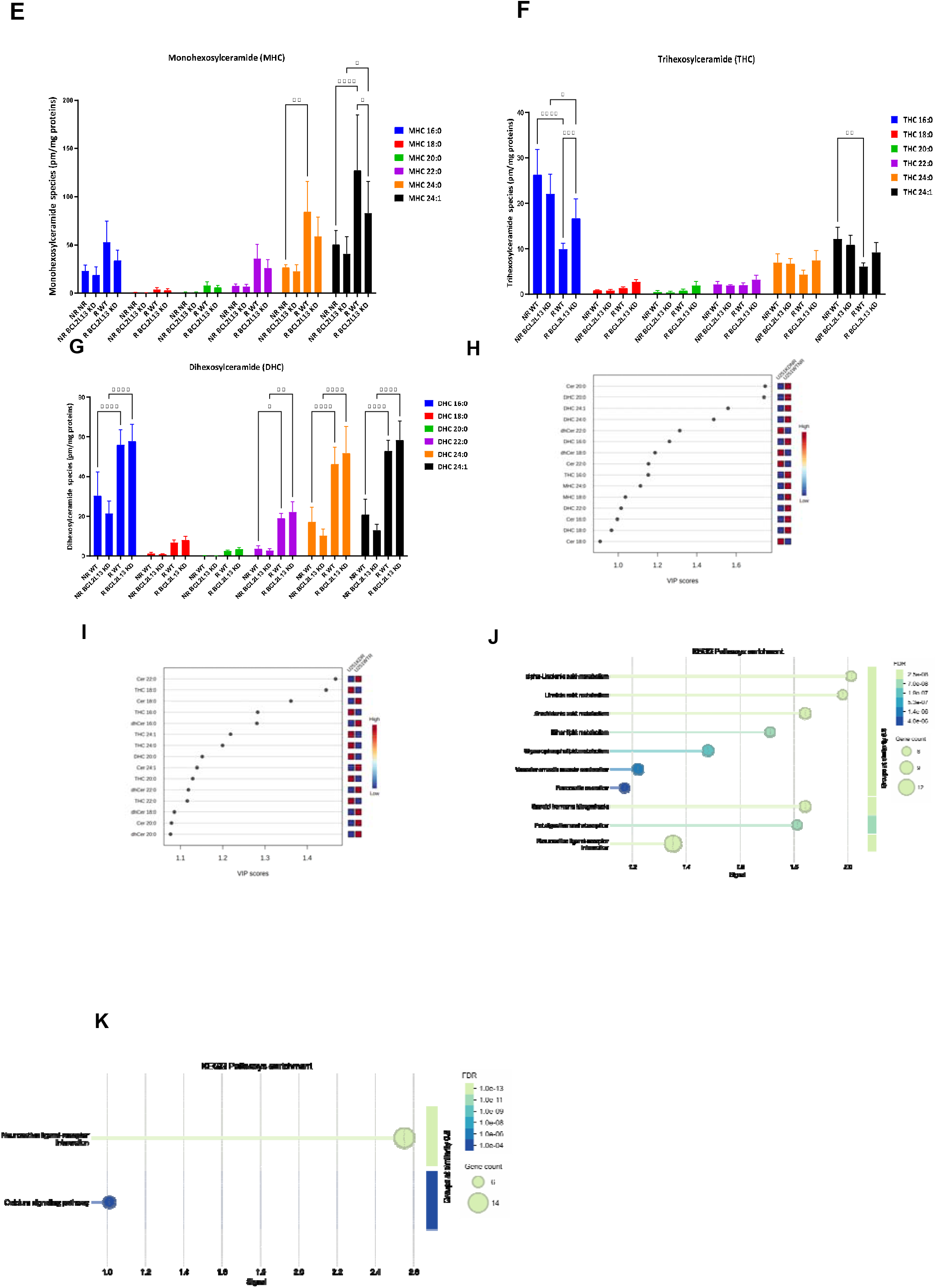
BCL2L13 Knockdown Rewires Ceramide Profiles via Resistance-Specific Deregulation of CerS2 and CerS6 Activity. Figure 6 shows the impact of BCL2L13 KD on ceramide metabolism in NR and R glioblastoma cells. (A) Heatmap analysis shows reduced levels of dihydroceramides (dhCer), monohexosylceramides (MHC), dihexosylceramides (DHC), and trihexosylceramides (THC) following KD, indicating loss of CerS2/CerS6 inhibition. (B) Principal component analysis (PCA) demonstrates distinct lipidomic separation between NR and R groups. (C–G) Quantification of ceramide subclasses reveals chain-length– and subclass-specific remodeling: Cer 16:0 and dhCer 16:0 increase in R KD; VLC dhCer, MHC, and DHC species are elevated in R WT; and THC species display diverse shifts depending on resistance state and glycosylation patterns. (H) Partial least squares discriminant analysis (PLS-DA) identifies Cer 20:0, DHC 20:0, 24:0, and 24:1 as key BCL2L13-regulated species in NR cells. (I) In R cells, Cer 22:0 and THC 18:0 are most discriminatory, aligning with CerS2 reactivation post-KD. (J, K) KEGG pathway enrichment reveals lipid metabolic shifts in NR KD cells and enhanced neuroactive and calcium signaling in R KD cells. All data represent n = 5 biological replicates.

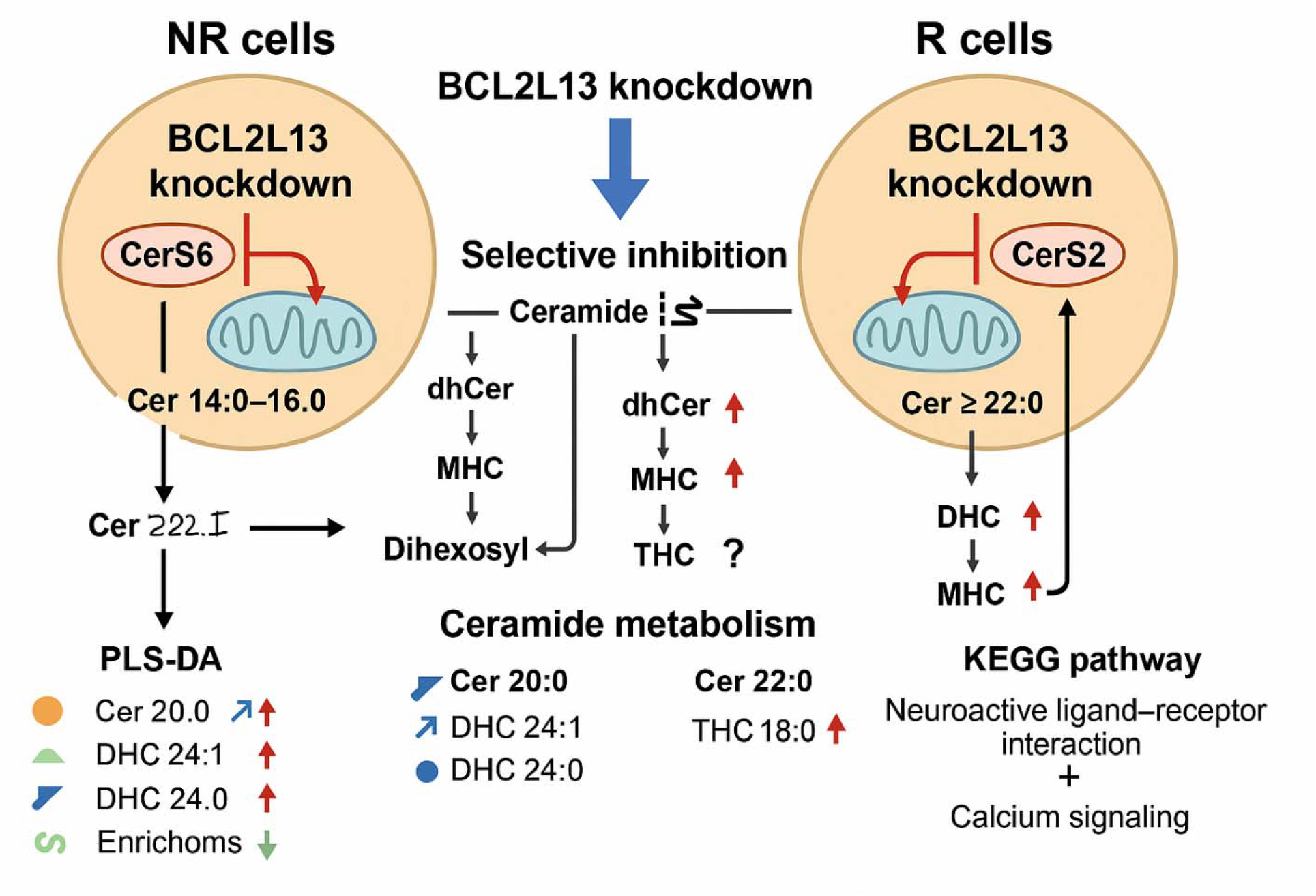

### Validation of KEGG Pathways Enriched Following BCL2L13 Knockdown in GBM Cells

Knockdown of BCL2L13 in NR cells led to significant enrichment of lipid metabolic pathways, including steroid hormone biosynthesis, arachidonic acid, linoleic acid, and α-linolenic acid metabolism (all FDR ≈2.5×10^–7) (Figure 6J). These findings suggest that loss of BCL2L13 perturbs lipid homeostasis, consistent with its role in autophagy and mitochondrial function. Notably, autophagy (particularly lipophagy) is essential for mobilizing cholesterol and fatty acids for steroidogenesis; inhibiting autophagy can dysregulate steroid hormone production and lipid metabolism gene expression [20]. BCL2L13 itself is a mitophagy receptor required for clearing damaged mitochondria [21]. Its deficiency leads to accumulation of dysfunctional mitochondria and oxidative stress, forcing cells to rewire their metabolism [21]. Accordingly, anti-apoptotic BCL-2 family proteins are known to support mitochondrial energetics beyond apoptosis – for example, BCL-2/BCL-X_L overexpression enhances oxidative phosphorylation and ATP generation in cancer cells (increasing Complex I activity and oxygen consumption [22]. Loss of BCL2L13 would thus be expected to compromise respiratory capacity and trigger compensatory upregulation of lipid biosynthetic pathways to meet energy demands. This aligns with our observation of increased LC3-II in BCL2L13-depleted sensitive cells (indicative of altered autophagic flux) (Figure 4) and the enrichment of lipid-signaling KEGG pathways, pointing to a metabolic stress response. Supporting this, Seahorse analysis (Figure 5) revealed that BCL2L13 knockdown in NR cells led to marked reductions in maximal and spare respiratory capacity and modestly impaired ATP-linked respiration, mimicking the low-capacity phenotype seen in resistant cells. These bioenergetic deficits may promote compensatory activation of lipid biosynthesis and remodeling pathways identified in KEGG enrichment.

In R cells, BCL2L13 knockdown primarily altered signaling pathways. KEGG analysis identified Neuroactive Ligand–Receptor Interaction as the top hit (14 genes, FDR ∼3.9×10^–13) and the Calcium Signaling pathway as second (6 genes, FDR ∼3.3×10^–4). These shifts highlight that disrupting BCL2L13 in the resistant context affects cell communication and Ca^2+^-homeostasis more than core metabolism. Notably, BCL-2 family proteins are key modulators of intracellular Ca^2+^ dynamics: BCL-2 at the ER inhibits IP3 receptor–mediated Ca^2+^ release, dampening calcium signaling and apoptotic cascades [23]. Thus, BCL2L13 loss may exacerbate Ca^2+^ dysregulation in TMZ-resistant cells, consistent with the enrichment of calcium-signaling genes (e.g. NOS2, HTR2C, TACR3). Furthermore, enrichment of the neuroactive ligand–receptor pathway (including opioid, cannabinoid, and serotonin receptors) suggests a broader stress-adaptive response in resistant GBM cells lacking BCL2L13. Such receptors have been implicated in glioma survival and may be upregulated as a compensatory mechanism when mitochondrial and autophagic function is impaired, echoing the known crosstalk between mitochondrial stress and cell signaling pathways [24]. Overall, our KEGG analysis validates that BCL2L13 depletion induces profound metabolic remodeling in sensitive cells (favoring lipid-related pathways) while provoking alterations in Ca^2+-linked signaling and receptor networks in resistant cells – results that are in line with BCL2L13’s established role in mitochondrial autophagy and the BCL-2 family’s influence on metabolism and calcium homeostasis. Furthermore, Seahorse metabolic profiling (Figure 5) demonstrated that BCL2L13 knockdown in TMZ-R cells, which already display profoundly reduced respiratory function, does not further impair maximal or ATP-linked respiration but exacerbates proton leak—highlighting persistent mitochondrial dysfunction potentially linked to calcium signaling dysregulation captured in KEGG analysis.

## DISCUSSION

TMZ-resistant glioblastoma cells (U251) exhibit a coordinated evasion of TMZ-induced apoptosis and a reprogramming of autophagy. In contrast to parental cells, the resistant cells maintain high viability even at 5-fold higher TMZ concentrations and show negligible sub-G0 apoptotic fractions, indicating suppression of apoptosis. This aligns with the elevated expression of BCL2L13 (Bcl-rambo) in resistant cells, as BCL2L13 is a Bcl-2 family member known to inhibit mitochondrial outer membrane permeabilization (MOMP) and caspase activation [14]. By preventing cytochrome *c* release, high BCL2L13 likely raises the threshold for TMZ-triggered apoptosis, helping resistant cells bypass the lethal DNA damage response. Notably, BCL2L13 is upregulated in high-grade and mesenchymal-subtype gliomas [11, 14, 25], consistent with its role in promoting tumor cell survival and invasiveness in GBM [11, 14, 25].

A striking feature of the TMZ-resistant cells is the disruption of autophagy flux. While TMZ-sensitive cells undergo autophagy (with LC3-II turnover and p62/SQSTM1 degradation), the resistant cells accumulate LC3-II and p62, signifying a block at the autophagosome–lysosome fusion step [5, 26, 27]. Indeed, adding the lysosomal inhibitor Bafilomycin A1 further increased LC3/p62 in sensitive cells but had no effect on resistant cells, confirming that autophagy was already maximally inhibited in the latter. BCL2L13 appears central to this phenotype. BCL2L13 localizes to mitochondria and functions as a mitophagy receptor by binding LC3 via a WXXI motif [21], driving the formation of autophagosomes containing damaged mitochondria. In resistant cells, high BCL2L13 may promote excessive mitophagosome formation without completion of degradation, leading to autophagosome accumulation. Consistently, BCL2L13 knockdown reversed this pattern: it reduced LC3-II levels (diminishing autophagosome buildup) in resistant cells while conversely causing LC3-II accumulation in non-resistant cells [28]. This inverse effect indicates that BCL2L13 helps set a balance between autophagosome formation and clearance depending on context. In the resistant state, BCL2L13-driven autophagy is trapped at an incomplete stage, which paradoxically might favor cell survival by preventing both autophagic cell death and excessive mitochondrial damage.

Mechanistically, the inhibition of autophagy flux in BCL2L13-high expression cells creates a cytoprotective milieu. One benefit is the stabilization of p62, an adaptor whose accumulation can activate the NRF2 antioxidant pathway [29]. Persistent p62 in the resistant cells likely sequesters Keap1, leading to NRF2-mediated induction of detoxification and survival genes, thereby mitigating TMZ-induced oxidative stress [30-32]. This autophagy/apoptosis coupling is reminiscent of canonical BCL-2:Beclin1 interactions in cancer cells, where Bcl-2 proteins concurrently block autophagy and apoptosis to optimize survival [33]. Similarly, BCL2L13 in our model serves as a dual regulator – it curtails apoptosis at the mitochondria while restraining autophagy to a non-lethal level. By removing damaged mitochondria via mitophagy yet halting their degradation, BCL2L13 helps resistant GBM cells avoid both energy crisis and the pro-death consequences of unchecked autophagy.

In a broader cancer context, these findings underscore how tumoral cells can co-opt BCL-2 family proteins to withstand therapy. Autophagy is often induced by chemo-radiotherapy as a stress tolerance mechanism [34, 35], and many tumors exhibit high basal autophagy to support growth or resist drugs. Notably, complete autophagy inhibition can be detrimental to cancer cells unless compensated by parallel adaptations [33, 36]. Here, GBM cells achieved such compensation: despite autophagy flux being blocked, the cells survived due to BCL2L13’s anti-apoptotic action and likely NRF2-driven stress relief. BCL2L13’s role may vary with cancer type (pro-tumor in GBM and leukemia, but less so in others [14]), yet its capacity to modulate both mitochondrial apoptosis and autophagy is a unifying theme. In essence, BCL2L13 acts as a molecular switch between life and death pathways – its upregulation in TMZ-resistant glioblastoma tilts the balance toward cell survival by inhibiting apoptosis and stalling autophagy flux, thereby preserving a state of cellular equilibrium under chemotherapeutic stress. This mechanistic insight into BCL2L13’s function provides a clearer understanding of how glioblastoma cells may evade TMZ and may inform future strategies to disrupt these survival circuits.

Based on these findings, our data highlight BCL2L13 as a critical nexus between mitochondrial integrity and ceramide metabolism in GBM. In TMZ-sensitive cells, BCL2L13 sustains oxidative metabolism – maintaining high respiratory capacity, ATP generation, and tight mitochondrial membranes – whereas TMZ-resistant cells have already lost much of this metabolic flexibility. Notably, BCL2L13’s context-dependent binding to ceramide synthases appears to underlie these differences. Consistent with earlier reports that BCL2L13 binds and inhibits CerS2 and CerS6 to curb proapoptotic ceramide production [11, 25], we observed that TMZ resistance alters this interaction landscape: BCL2L13 preferentially associates with CerS6 in sensitive cells but with CerS2 in resistant cells. Consequently, BCL2L13 knockdown unleashes CerS activity in a state-specific manner, elevating the respective ceramide species and their metabolites. For example, loss of BCL2L13 markedly increased C16:0 ceramide (a CerS6 product) and related glycosylated ceramides in the sensitive background, while in resistant cells it boosted very-long-chain CerS2 outputs like C22:0 ceramide and downstream di- and tri-hexosylceramides. These lipidomic shifts – corroborated by PCA/PLS-DA – indicate that BCL2L13 actively restrains ceramide accrual and rerouting. The enrichment of specific lipid pathways (e.g. steroid biosynthesis in sensitive cells, calcium signaling in resistant cells) further underscores the broad metabolic repercussions of disrupting BCL2L13. Mechanistically, this aligns with evidence that TMZ-resistant gliomas evade ceramide-driven cell death by metabolizing ceramides into less toxic forms [37]. Overall, BCL2L13 preserves mitochondrial function and limits pro-death sphingolipid accumulation in GBM, a mechanistic adaptation that not only fortifies TMZ-sensitive cells but is co-opted in resistance. While our focus is GBM, the dynamic BCL2L13–CerS axis may exemplify a wider cancer strategy to rewire lipid metabolism for survival.

Our study defines BCL2L13 as a central node linking mitochondrial homeostasis, autophagy regulation, and lipid metabolism in glioblastoma. In TMZ-resistant cells, elevated BCL2L13 suppresses mitochondrial apoptosis and traps autophagy at an incomplete stage, generating a cytoprotective state that sustains survival under genotoxic stress. Conversely, in TMZ-sensitive cells, BCL2L13 maintains oxidative capacity and limits ceramide-driven stress by selectively associating with CerS6, whereas resistance is accompanied by a shift toward CerS2 coupling and altered lipid remodeling. The integration of LC3/p62 flux assays, Seahorse bioenergetics, and lipidomic–KEGG pathway analyses internally validates that BCL2L13 orchestrates a metabolic–autophagic network that buffers mitochondrial and lipid stress.

While BCL2L13 knockdown does not resensitize resistant cells to TMZ, it reveals how tumor cells reprogram bioenergetics and lipid homeostasis to sustain growth. Limitations include the use of a single in vitro lineage and absence of genetic rescue and in vivo validation. Future studies should investigate the BCL2L13–CerS axis in patient-derived and orthotopic GBM models, assess correlations with clinical outcome, and test whether targeting BCL2L13-dependent mitophagy or lipid pathways can disrupt metabolic resilience and improve therapeutic response in glioblastoma.

## DATA AVAILABILITY

Data will be available by corresponding author.

## ACKNOWLEDGEMENTS

The authors acknowledge CHATGPT 4o for professional English edit.

## AUTHOR CONTRIBUTIONS

Courtney Clark (C.C.) conducted all experimental work and prepared the methods section. Amir Barzegar-Behrooz (A.B.B.) and Mehdi Eshraghi (M.E) drafted the manuscript and performed the literature review. Simone C. da Silva Rosa (S.C.S.R.), Jaodi Jacobs (J.J.), Kianoosh Naghibzadeh (K.N), and Xiaohui Wang (X.W.) conducted the lipidomics analysis and prepared the related figures. Abhay Srivastava (A.S.) performed the Seahorse assays and analyzed the data. Rui Vitorino (R.V.) supervised the lipidomics analysis and contributed to drafting the lipidomics section. Sudharsan Rao Ande (S.R.A.) established and executed the Co-immunoprecipitation (Co-IP) experiments. Amir Ravandi (A.R.) led the lipidomics experimental planning. Sanjiv Dhingra (S.D.) provided supervision and leadership in mitochondrial metabolism analysis. Stevan Pecic (S.P.) contributed to supervision, resources, and initial manuscript drafting. Donal Miller (D.M.) provided supervision, resources, and conceptualization of the study. Shahla Shojaei (S.S.) developed the Resistant cells, and contributed to project design and conceptualization. Saeid Ghavami (S.G.) led and supervised the project, oversaw study design, and secured resources.

## ADDITIONAL INFORMATION

N/A

